# Methods to study toxic transgenes in *C. elegans*: an analysis of protease-dead separase in the *C. elegans* embryo

**DOI:** 10.1101/002444

**Authors:** Diana M. Mitchell, Lindsey R. Uehlein, Joshua N. Bembenek

## Abstract

We investigated whether the protease activity of separase, which is required for chromosome segregation, is also required for its other roles during anaphase in *C. elegans* given that non-proteolytic functions of separase have been identified in other organisms. We find that expression of protease-dead separase is dominant-negative in *C. elegans* embryos. The *C. elegans* embryo is an ideal system to study developmental processes in a genetically tractable system. However, a major limitation is the lack of an inducible gene expression system for the embryo. The most common method for embryonic expression involves generation of integrated transgenes under the control of the *pie-1* promoter, using *unc-119* as a selection marker. However expression of dominant-negative proteins kills the strain preventing analysis of mutants. We have developed two methods that allow for the propagation of lines carrying dominant-negative transgenes in order to study protease-dead separase in embryos. The first involves feeding *gfp* RNAi to eliminate transgene expression and allows propagation of transgenic lines indefinitely. Animals removed from *gfp* RNAi for several generations recover transgene expression and associated phenotypes. The second involves propagation of the transgene with the female specific *pie-1* promoter via the male germline and analysis of phenotypes in embryos from F1 heterozygous hermaphrodites that express the protein. Using these methods, we show that protease-dead separase causes chromosome nondisjunction and cytokinesis failures. These methods are immediately applicable for studies of dominant-negative transgenes and should open new lines of investigation in the *C. elegans* embryo.

## INTRODUCTION

Experimental design to discover protein functions often involves expression of mutant form of a protein of interest *in vivo*, which can interrupt cellular processes and shed light on protein function, but also causes lethality. In several systems, methods have been developed to allow for inducible expression of such mutant proteins to facilitate their study. In the *C. elegans* embryo, the method for expression of fusion proteins involves generation of integrated transgenes under the control of an oocyte promoter and maternal deposition of these transgenic proteins. However, this system is not inducible and prevents generation of lines expressing dominant-negative or otherwise “toxic” proteins because worm strains are inviable. Methods are needed to allow inducible expression in order to study proteins that cause lethality in this system.

Separase is a cysteine protease with multiple roles during cell division. In a number of these roles the protease activity of separase is required, including cohesion cleavage at the onset of anaphase (Uhlmann *et al.* 1999; Uhlmann *et al.* 2000; Waizenegger *et al.* 2000; Hauf *et al.* 2001), DNA repair (Nagao *et al.* 2004), resolution of chiasmata in mouse oocytes (Kudo *et al.* 2006), mitotic spindle elongation (Baskerville *et al.* 2008), and centriole duplication (Schockel *et al.* 2011; Matsuo *et al.* 2012). Additional non-proteolytic functions of separase have been identified, including anaphase exit (Sullivan and Uhlmann 2003) and Cdc14 early anaphase release (FEAR) pathway activation (Ross and Cohen-Fix 2004; Lu and Cross 2009), and polar body extrusion in mouse oocytes (Kudo *et al.* 2006). We have demonstrated that separase is crucial for proper membrane trafficking in the *C. elegans* embryo. Separase is required for exocytosis of cortical granules following fertilization (Bembenek *et al.* 2007) that ensure proper formation of the embryonic eggshell. Separase is also required for cytokinesis and the regulation of RAB-11 vesicles at the cleavage furrow and midbody following mitosis (Bembenek *et al.* 2010). Another report demonstrated that separase also regulates membrane trafficking during cytokinesis in *Arabidopsis* (Moschou *et al.* 2013), and separase is associated with membranes in human cells (Bacac *et al.* 2011) suggesting that separase’s membrane trafficking function is conserved among organisms. However, the mechanism behind separase’s membrane trafficking role is not known.

A key question is whether the protease activity of separase is required for exocytosis. We created a transgenic worm line with a protease-dead form of separase fused to GFP (SEP-1^PD^::GFP) using standard methods to address this question ((Bembenek *et al.* 2010), and Figure 1). Unexpectedly, SEP-1^PD^::GFP expression caused embryo lethality, causing the line to die out within a few generations under normal growth conditions. Other researchers have encountered similar difficulties with other mutant proteins (Hao *et al.* 2006), highlighting a need for methods to control expression of toxic transgenes in the worm embryo. Accordingly, we have developed methods to maintain transgenic worm lines that carry “toxic” transgenes and allow for subsequent expression of the transgene for experimental and genetic analysis of it’s function.

**Figure 1.**
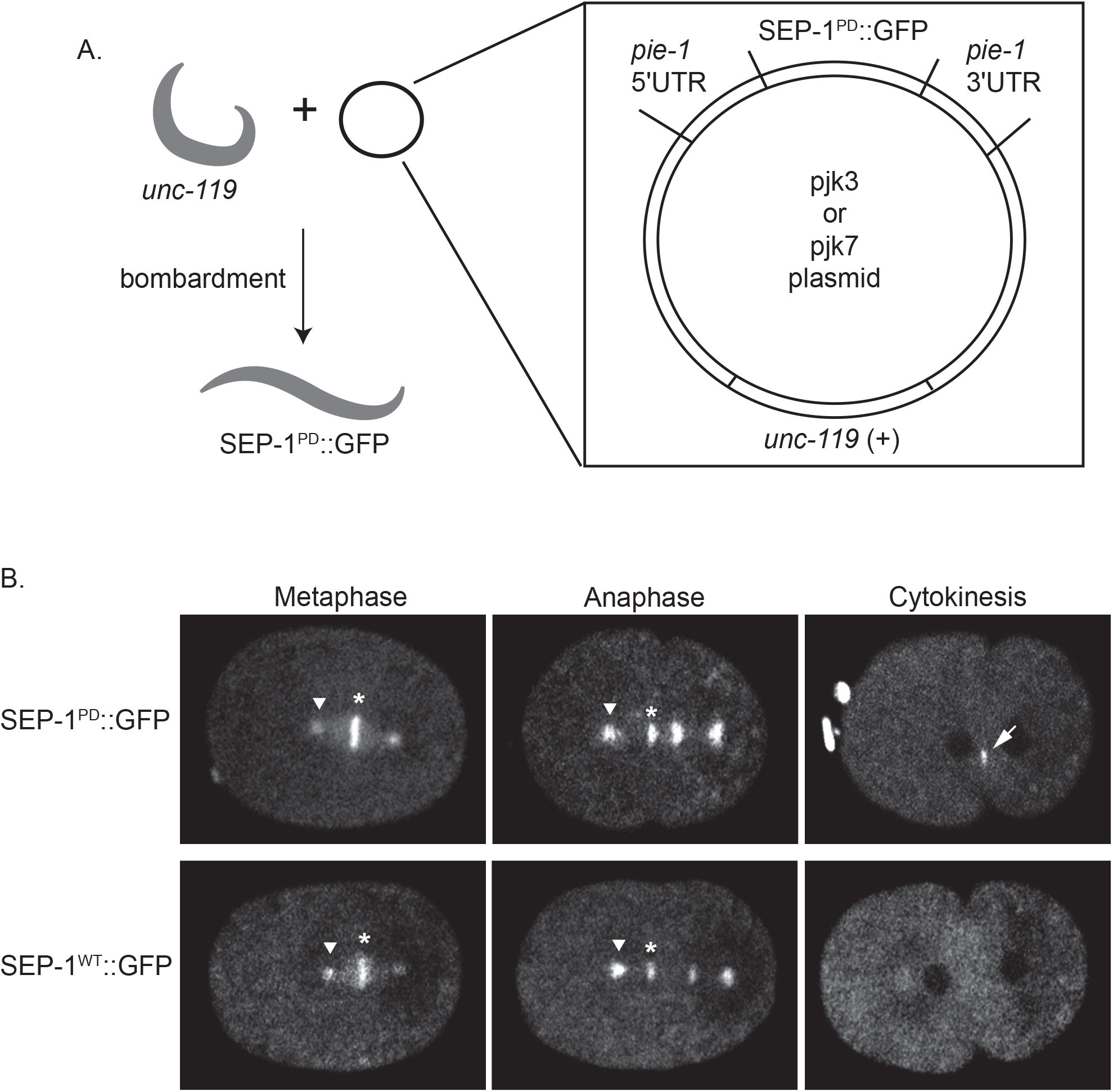
Creation of SEP-1^PD^::GFP transgenic worm lines. A. Microparticle bombardment of *unc-119* worms with plasmid DNA bound to gold beads. The plasmid (enlarged panel) contains the *sep-1* coding sequence with mutation in the protease domain fused to GFP under control of the *pie-1* promoter and an *unc-119* rescue sequence, allowing for identification of transformed worm lines. B. SEP-1^PD^::GFP (top row) and SEP-1^WT^::GFP (bottom row) localization in embryos during the indicated stages of mitosis. SEP-1^WT^::GFP and SEP-1^PD^::GFP localizes to chromosomes (star) and centrosomes (arrowhead). During cytokinesis, SEP-1^PD^::GFP accumulates at the cleavage furrow (arrow).

The first method involves feeding transgenic worms *gfp* RNAi, which silences transgene expression, and allows for maintenance of homozygous transgenic lines indefinitely. Upon removal from *gfp* RNAi, transgene re-expression accompanies the return of associated phenotypes. The second method uses male worms to propagate the transgene. Because SEP-1^PD^::GFP expression is under control of the *pie-1* promoter, sperm do not express the toxic fusion protein (Mello *et al.* 1996) and transgenic male worms are viable. Transgenic males can be crossed to *unc-119* hermaphrodites to generate transgenic hermaphrodite progeny for study, as well as crossed to mutant and other transgenic lines for further analysis. These methods employ standard laboratory techniques used by *C. elegans* researchers, which will open new possibilities for analysis of gene function in the *C. elegans* embryo.

We have successfully used these newly developed methods to characterize SEP-1^PD^::GFP in the *C. elegans* embryo. Embryos expressing SEP-1^PD^::GFP showed high levels of embryonic lethality, even in the wild-type background. We conclude that protease-dead separase is dominant negative and interferes with endogenous separase function. In further support of such dominant negative activity, we found that SEP-1^PD^::GFP generates exacerbated phenotypes when expressed in heterozygous separase mutants. Further, SEP-1^PD^::GFP expressing embryos had slower chromosome separation during mitotic anaphase, as well as frequent lagging chromosomes and chromosome bridges, compared to embryos expressing SEP-1^WT^::GFP. Moreover, SEP-1^PD^::GFP embryos displayed multipolar spindles after cytokinesis failure during mitosis. These results suggest that separase promotes cytokinesis by a proteolytic mechanism in *C. elegans*.

## MATERIALS AND METHODS

### Strains

*C. elegans* strains were maintained according to standard protocols (Brenner, 1974). Temperature sensitive strains were maintained at 16°C, unless otherwise indicated in the text, and shifted to non-permissive temperature as indicated. All other strains were maintained at 20°C. Strains containing the protease-dead *sep-1* transgene were maintained on lawns of *gfp* RNAi feeding bacteria as indicated in text and below, then transferred onto OP-50 lawns as indicated.

Strains used in this study are indicated in the following table:

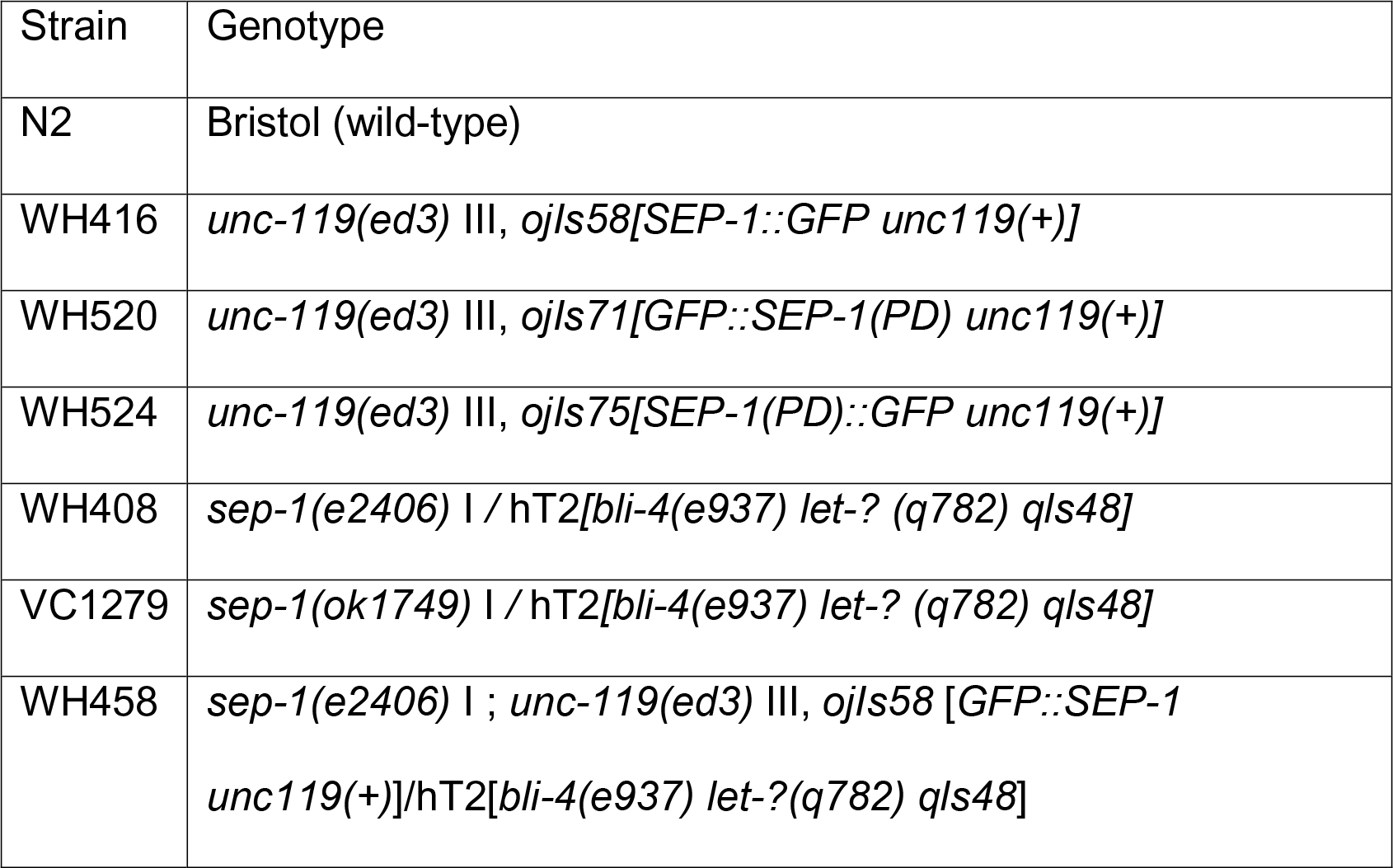

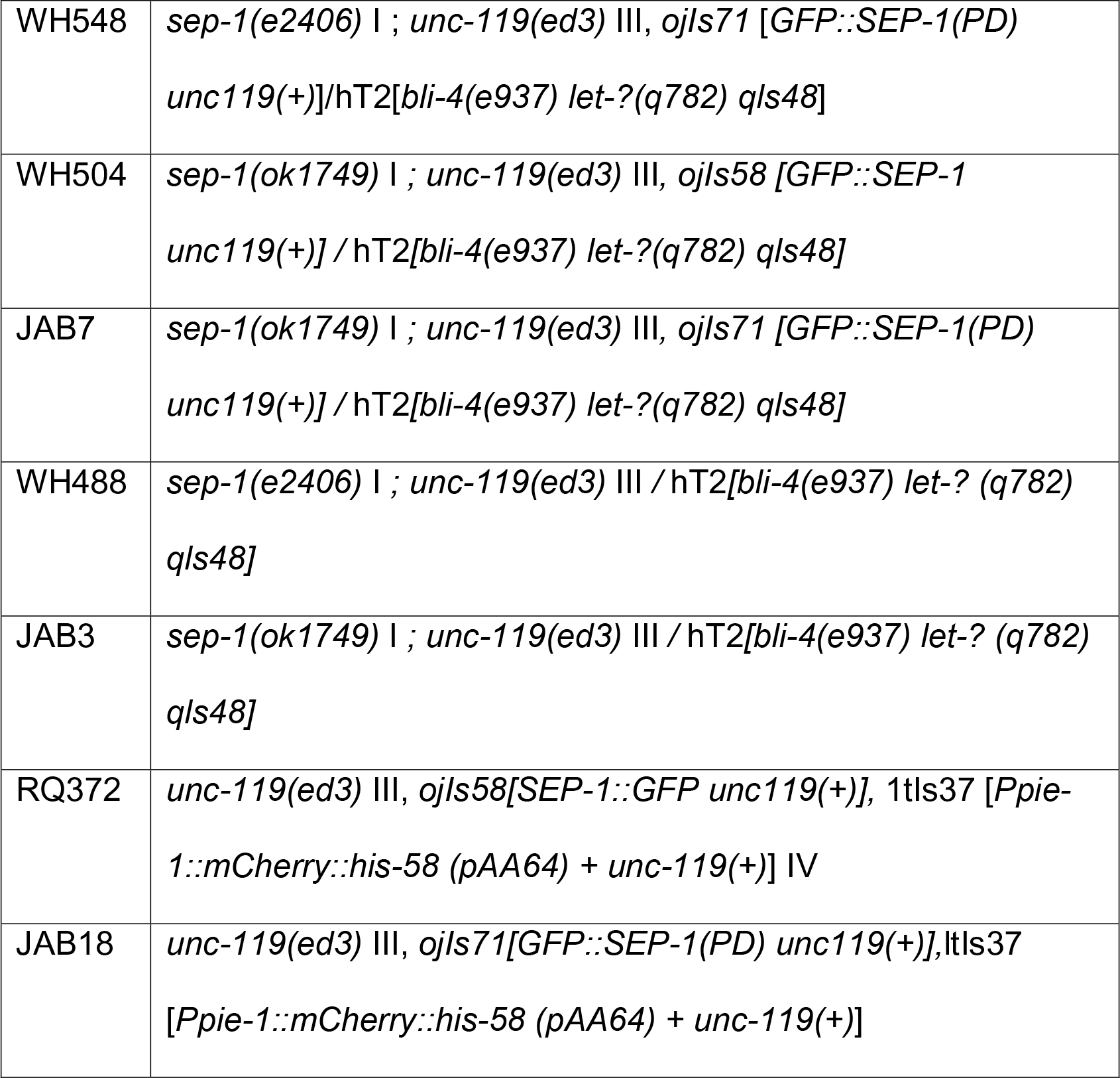

JAB18 was obtained by crossing WH520 males with OD56 hermaphrodites, and subsequent generations were maintained on *gfp* RNAi. At each generation following the cross, approximately half of the worms at L4 stage were moved to OP50 plates for 24 hours at 25°C and screened for the presence of both transgenes by microscopy. Worms were then singled from the original *gfp* RNAi feeding plate. This protocol was repeated until double homozygous transgenic lines were obtained, after which the line was maintained on *gfp* RNAi at 20°C. Feeding *gfp* RNAi did not silence expression of H2B::mCherry (data not shown).

### Molecular Biology

Cloning the protease-dead separase mutant DNA sequence into pjk3 or pjk7 vectors was performed as previously described (Bembenek *et al.* 2010). Microparticle bombardment (Praitis *et al.* 2001) was used to obtain transgenic worm lines as described in the text and Figure 1.

### RNAi feeding

The *gfp* RNAi feeding construct in L440 vector was obtained from Dr. Scott Kennedy. To silence GFP fusion transgenes and maintain worm lines, worms were picked onto lawns of *gfp* RNAi feeding bacteria and L4 worms were picked at each generation onto fresh lawns. In order to provide the optimal RNAi effect for transgene silencing, worms were grown on *gfp* RNAi at 20°C (which is the semi-permissive temperature for the *sep-1(e2406)* allele), as we were unable to propagate some lines on *gfp* RNAi by feeding at 16°C. For transgene re-expression, L4 worms were removed to OP50 lawns and picked onto fresh OP50 feeding plates at each generation as indicated in the text and figures.

### Eggshell Permeability

L4 hermaphrodites were singled onto NGM plates containing (150 μg/mL) Nile Blue dye (Sigma-Aldrich) seeded with a spot of OP50. After 24 hours, mothers were removed and the number of blue (permeable) and non-blue (non-permeable) embryos was counted, then the percentage of permeable embryos was determined.

### Microscopy

For imaging of mitotic embryos, young adult worms were dissected in M9 and embryos were mounted on agar pads as previously described (Bembenek *et al.* 2010). For imaging of meiotic embryos, hanging drop method was used as previously described (Bembenek *et al.* 2007). Live cell imaging was performed using a Nikon Eclipse CSU-22 inverted confocal spinning disc imaging system equipped with a 60X 1.40NA objective from Visitech International, running metamorph software. Digital images were obtained with a Photometrics EM-CCD camera. Movies were analyzed using FIJI (ImageJ) software using the Bio-Formats plugin from LOCI (www.loci.wisc.edu).

## RESULTS

### Creation of protease-dead separase transgenic worm lines

We created SEP-1^PD^::GFP expressing transgenic worm lines using microparticle bombardment, using standard protocols (Praitis *et al.* 2001), but GFP expressing lines could not be maintained for more than a few generations. The final construct contains several features that allowed us to propagate lines carrying dominant negative transgenes. The construct is designed to generate proteins fused to GFP driven by the *pie-1* promoter (Figure 1A). Importantly, the *pie-1* promoter is widely used to drive transgene expression in *C. elegans* embryos and is strongly biased toward oocyte (maternal) expression (Mello *et al.* 1996). The construct also has an *unc-119* rescue sequence allowing for identification of transformed worms. We cloned genomic *sep-1* sequence, with a point mutation that results in Cysteine to Serine substitution at amino acid 1040, located in the protease domain of SEP-1. We created multiple independent worm lines with integrated transgenes coding for both N- and C-terminal fusions of GFP to SEP-1^PD^ using this strategy, all of which had severe growth defects.

Examination of SEP-1^PD^::GFP expressing embryos showed that SEP-1^PD^::GFP localizes to putative sites of separase activity similarly to SEP-1^WT^::GFP during mitosis (Figure 1B and previously reported (Bembenek *et al.* 2010)). SEP-1^PD^::GFP accumulates more strongly than SEP-1^WT^::GFP at several sites of action during anaphase of mitosis, including the cleavage furrow during cytokinesis (Figure 1B and previously reported (Bembenek *et al.* 2010)). These sites of action may have substrates of separase, which may have stronger binding to the inactive protease, suggesting that SEP-1^PD^::GFP could be substrate trapping. If protease-dead separase remains bound to substrates, it could interfere with their cleavage by endogenous separase having a dominant-negative effect. Consistent with a dominant negative activity, SEP-1^PD^::GFP expression caused embryo lethality (Figure 2 and Supplemental Figure 1, see below) in the wild-type background with two copies of normal endogenous separase. Dominant negative activity of protease-dead separase has not been reported in other systems. High levels of embryonic lethality in SEP-1^PD^::GFP expressing worm lines required us to develop methods to propagate this “toxic” transgene in order to further examine the SEP-1^PD^::GFP.

**Figure 2.**
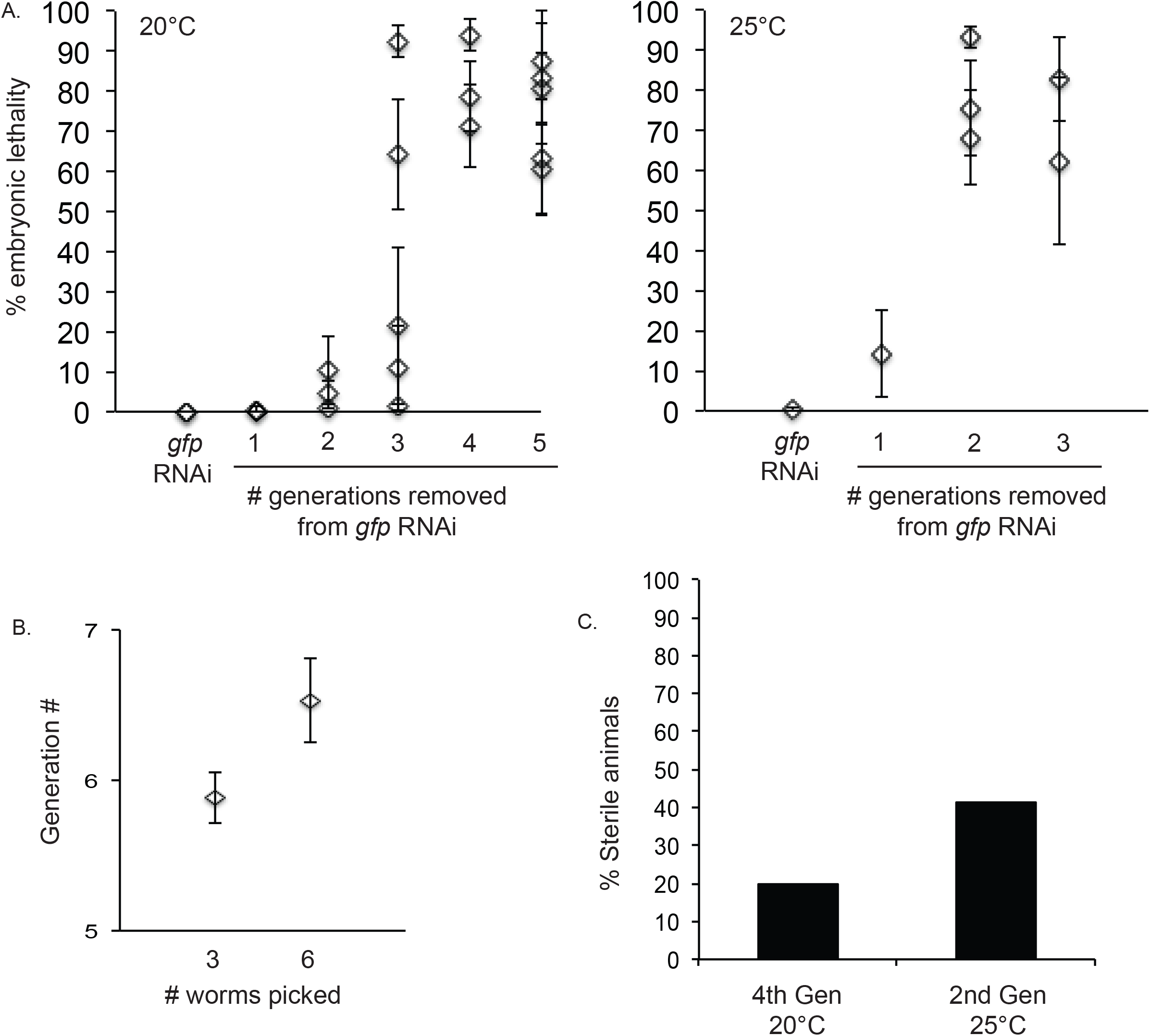
SEP-1^PD^::GFP worm lines can be maintained on *gfp* RNAi. A. Embryonic lethality of WH520 (SEP-1^PD^::GFP) line on gfp RNAi and after removal onto OP50 plates at 20°C (left) or 25°C (right). Each data point with error bars represents the average of a group of 10 singled worms +/− SEM examined in an individual experiment. B. Average generation +/− SEM that could be propagated for the WH520 line after removal from *gfp* RNAi when the indicated number of worms are picked at each generation and kept at 20°C. C. Percentage of sterile animals in the WH520 line after removal from *gfp* RNAi.

### Toxic transgene propagation method 1: Silencing of GFP expression by RNAi

RNAi provides a reliable system for targeted gene knock down in *C. elegans*. We took advantage of RNAi in order to silence expression of the SEP-1^PD^::GFP transgene by maintaining transgenic worm lines on lawns of *gfp* RNAi feeding bacteria. After bombardment following the standard protocol, UNC-rescued animals were screened for GFP expression, and non-Unc GFP positive lines had high lethality when grown under standard lab conditions. However, SEP-1^PD^::GFP transgenic worm lines fed *gfp* RNAi showed no embryonic lethality and could be propagated indefinitely at 20°C and 25°C (Figure 2A). When worms were transferred from *gfp* RNAi onto OP50, embryonic lethality returned after several generations (Figure 2A), and higher levels of embryonic lethality correlated with higher GFP expression levels (data not shown). The return of embryonic lethality was dependent on temperature, as embryonic lethality occurred in 3-5 generations at 20°C and 2-3 generations at 25°C, which could be due to reduced generational *gfp*(RNAi) transmission, increased transgene expression at 25°C, or an increase in cell cycle timing leading to a decrease in fidelity of division. Picking a larger number of worms at each generation allows for propagation of the transgenic line on OP50 through one more generation (Figure 2B). This could be due to affects with picking animals of different penetrance of generational RNAi propagation, as seen previously (Alcazar *et al.* 2008). Similar results were obtained for multiple independent worm lines expressing both N-terminal and C-terminal SEP-1^PD^ GFP fusion proteins, indicating that the position of GFP fusion is not a factor (Supplemental Figure 1). SEP-1^PD^ expressing worms that survive hatching show abnormal developmental phenotypes including tail defects, slow growth, and sterility (Figure 2C and data not shown), suggesting that protease-dead separase interferes with normal developmental processes in addition to causing embryonic lethality. Because transgene expression is *pie-1* driven, and should be most highly expressed in the germline and deposited in the egg, these results suggest that SEP-1^PD^ expressing worms show phenotypes that are a result of defects in the developing embryo.

### Toxic transgene propagation method 2: Male propagation

Expression of most transgenes in the *C. elegans* embryo, including our SEP-1^PD^::GFP transgene, is under control of the *pie-1* promoter, which is oocyte specific and is not expressed in sperm (Mello *et al.* 1996). We therefore reasoned that we could propagate the SEP-1^PD^::GFP transgene in the male germline where transgene expression is expected to be limited. We created SEP-1^PD^::GFP male worms by heat shock and backcrossed to *unc-119* hermaphrodites (Figure 3A). In the F1 progeny, heterozygous transgenic males can be used for propagation of the transgene and resulting hermaphrodites for transgene analysis. Transgenic heterozygous males were continually backcrossed to *unc-119* hermaphrodites, and were also used in crosses with other worm lines to test for genetic interactions. Single F1 heterozygous transgenic SEP-1^PD^::GFP hermaphrodites were picked onto individual plates and their progeny were analyzed for embryonic lethality. We found that embryonic lethality in the F2 brood was consistently in the range of 40-60% (Figure 3B). Even after backcrossing males to *unc-119* hermaphrodites more than 50 generations, embryonic lethality in the F2 remained within this range (Figure 3C). Therefore, propagation of *C. elegans “*toxic” *pie-1* driven transgenes in males bypasses lethality and can be done indefinitely, while also providing consistent transgene expression in the F1 generation.

**Figure 3.**
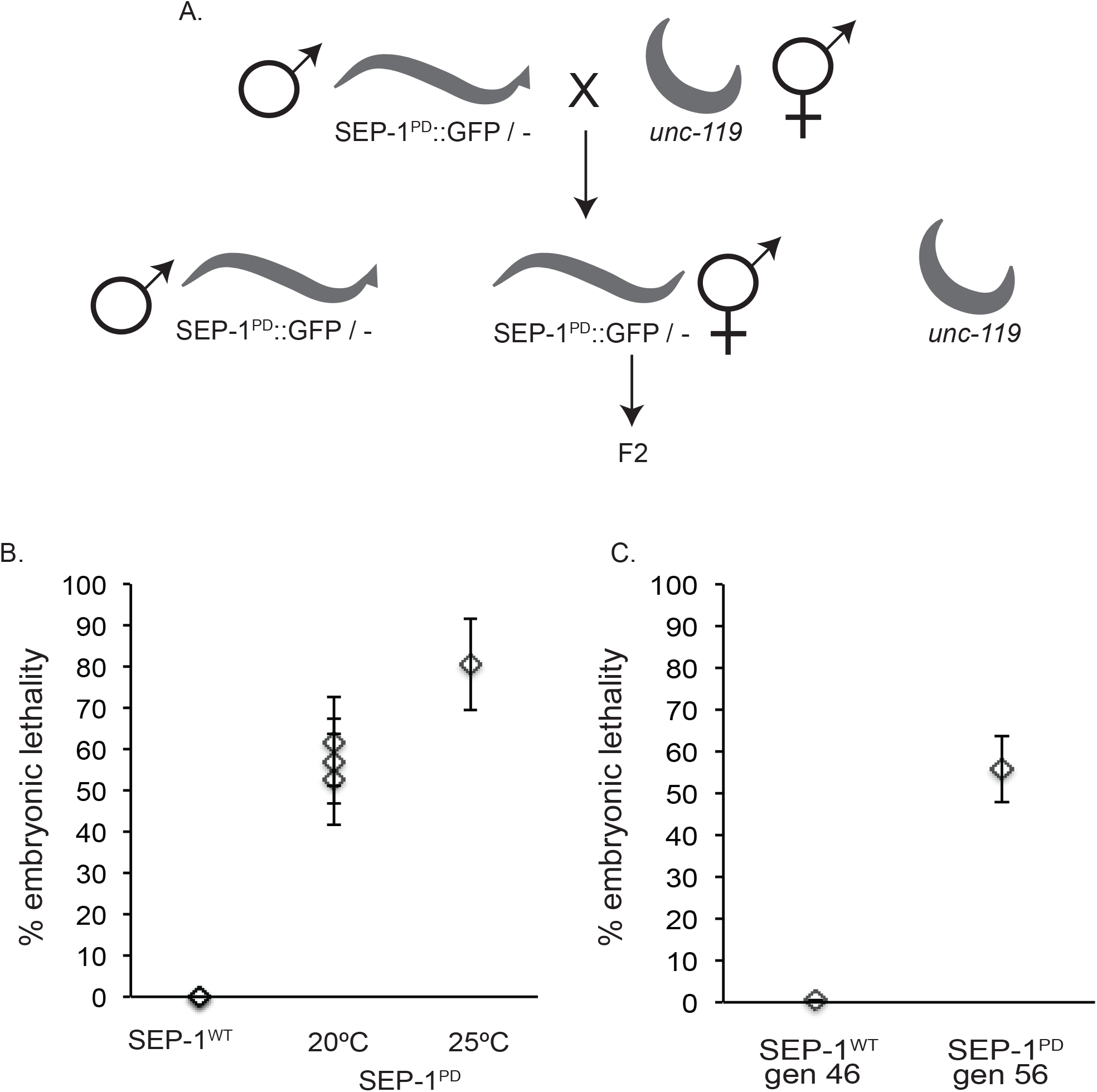
Propagation of the SEP-1^PD^::GFP transgene by backcrossing. A. The diagram represents the strategy used to propagate the SEP-1^PD^::GFP transgene using males. Because the transgene is under control of the *pie-1* promoter, expression in sperm is not significant. SEP-1^PD^::GFP/− transgenic males are continually crossed to *unc-119* hermaphrodites to generate heterozygous SEP-1^PD^::GFP/− males and hermaphrodites. B. Embryonic lethality in F2 from singled F1 heterozygous SEP-1^WT^::GFP/− or SEP-1^PD^::GFP/− hermaphrodites (generated by the male propagation method shown in A) at the indicated temperature. C. Embryonic lethality in the F2 after the indicated number of backcrosses of heterozygous SEP-1^WT^::GFP/− or SEP-1^PD^::GFP/− transgenic males to *unc-119* hermaphrodites. Data points represent the average of a group of 10 singled worms +/− SEM.

### Genetic interactions of protease-dead separase with separase mutants

Since transgenic SEP-1^PD^::GFP worms also express endogenous wild-type separase, we examined genetic interactions of SEP-1^PD^::GFP with hypomorphic *sep-1(e2406)* and *sep-1(ok1749)* deletion mutants. The *SEP-1(e2406)* protein contains an intact protease domain, but cannot localize to vesicles (Bembenek *et al.* 2007). We therefore examined embryos expressing SEP-1^WT^::GFP or SEP-1^PD^::GFP, which can localize to vesicles in meiosis I (not shown), in *sep-1(e2046)* mutants. The *sep-1(e2406)* mutant is temperature sensitive and viable at 16°C, but only viable as a balanced heterozygote at 20°C. The *sep-1(ok1749)* deletion mutant is likely a null allele since no protein can be detected by western blot (Richie *et al.* 2011). At all temperatures, nearly all homozygous *sep-1(ok1749)* progeny die during embryogenesis, with very few surviving animals that arrest at early larval stages. Both *sep-1(e2406)* and *sep-1(ok1749)* are maintained as balanced heterozygotes with the *hT2*[*bli-4(e937) let-? (q782) qls48*] balancer chromosome which encodes GFP localized to the pharynx (hT2 g).

We generated lines that were homozygous for either the SEP-1^WT^::GFP or SEP-1^PD^::GFP transgenes in these balanced separase mutant backgrounds. Both the balanced heterozygous *sep-1(e2406)* and *sep-1(ok1749)* deletion mutants were extremely sick when crossed with the SEP-1^PD^::GFP transgene and could not be maintained on normal OP50 bacterial plates. Therefore, the balanced separase mutant lines with SEP-1^PD^::GFP were maintained on *gfp* RNAi, yet even under this condition the strains are sick compared to SEP-1^WT^::GFP expressing lines. In addition to reduced viability, the progeny of the balanced *sep-1(e2406)* mutant animals expressing SEP-1^PD^::GFP produces few surviving homozygous mutant animals at 16°C (data not shown). In order to provide the optimal RNAi effect for transgene silencing, worms were grown on *gfp* RNAi at 20°C (which is the semi-permissive temperature for the *sep-1(e2406)* allele), as we were unable to propagate these lines on *gfp* RNAi by feeding at 16°C. Furthermore, the balanced separase mutant lines homozygous for SEP-1^PD^::GFP could only be propagated for a maximum of 1-3 generations off of *gfp* RNAi at 20°C before all progeny died, arrested prior to reaching adult, or were completely sterile (data not shown). This data indicates that SEP-1^PD^::GFP produces more severe phenotypes in a separase loss-of-function background.

We analyzed lethality in embryos from heterozygous, balanced mutant animals homozygous for SEP-1^WT^::GFP or SEP-1^PD^::GFP transgenes (Figure 4A). We examined embryonic lethality in the balanced animals in the third generation removed from *gfp* RNAi. As expected, we found that embryonic lethality in balanced mutant lines homozygous for SEP-1^PD^::GFP was more severe than the mutant alone, and this effect was reduced when transgene expression was silenced with *gfp* RNAi (Figure 4B and 4D). Further, SEP-1^WT^::GFP expression was able to rescue both homozygous *sep-1(e2406)* and *sep-1(ok1749*) mutant progeny while SEP-1^PD^::GFP could not (Figure 4C and 4E). Importantly, expression of SEP-1^WT^::GFP can rescue both homozygous *sep-1(e2406)* hypomorphic and *sep-1(ok1749)* deletion mutants to produce a few gravid adult animals (not shown). Unexpectedly, we saw slightly higher levels of embryonic lethality in broods from *sep-1(e2406 or ok1749)*/hT2g; SEP-1^WT^::GFP animals, even though SEP-1^WT^::GFP rescued the unbalanced mutant homozygous progeny (Figure 4B and 4D). This might reflect that SEP-1^WT^::GFP is not fully active and could also interfere with separase activity, which does not lead to phenotypes in WT cells, but is more prominent in mutant backgrounds. In addition, it was previously observed that protein dosage levels at various intracellular locations impacts separase function (Schvarzstein *et al.* 2013). Due to this stringent subcellular localization, SEP-1^WT^::GFP might cause an imbalance in separase localization and activity that causes lethality and leads to more prominent phenotypes in partial loss of function backgrounds. Consistent with this hypothesis, feeding *sep-1(e2406* or *ok1749)*/hT2g;SEP-1^WT^::GFP animals *gfp* RNAi restored levels of embryonic lethality and the distribution of balanced and unbalanced progeny back to the levels observed in *sep-1(e2406 or ok1749*)/hT2g without the transgene (Figure 4B–D). These data indicate that protease-dead separase does not rescue viability in separase mutant embryos.

**Figure 4.**
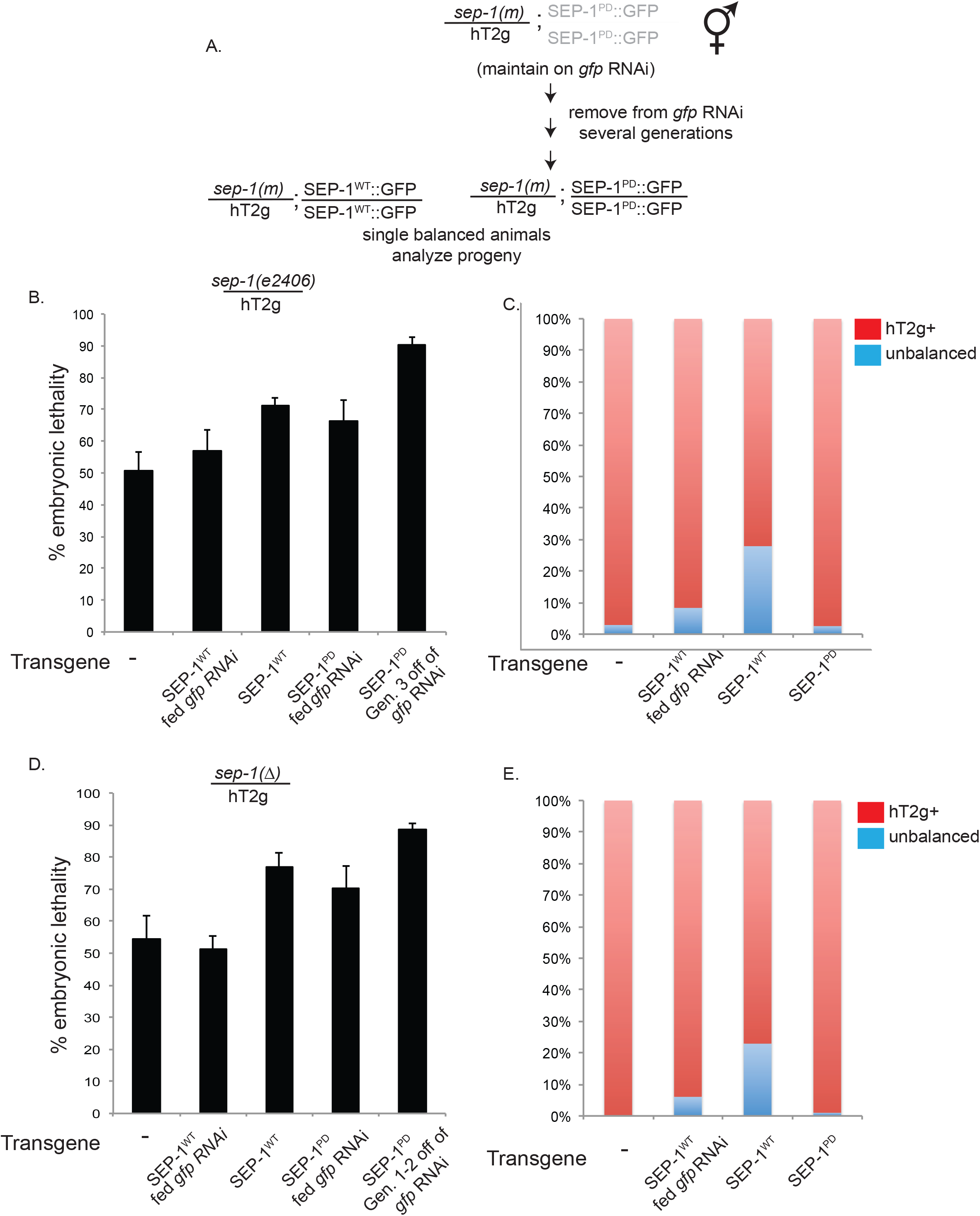
Genetic interactions of wild-type and protease-dead SEP-1 with *sep-1* mutant alleles in balanced mutant lines with homozygous transgenes. A. Lines with SEP-1^PD^::GFP transgene were maintained on *gfp* RNAi at 20°C and removed for several generations to allow transgene expression. Balanced worms were then singled and progeny were analyzed. B. Embryonic lethality in brood from singled *sep-1(e2406)*/hT2g hermaphrodites with homozygous SEP-1^WT^::GFP or SEP-1^PD^::GFP transgene. C. Percentage of progeny that are balanced mutant or homozygous mutant from singled *sep-1(e2406*)/hT2g hermaphrodites homozygous for the indicated transgene. D. Embryonic lethality in brood from singled *sep-1(ok1749*)/hT2g (referred to as Δ) hermaphrodites homozygous for the SEP-1^WT^::GFP or SEP-1^PD^::GFP transgene at 20°C. E. Percentage of progeny that are balanced mutant or homozygous mutant from singled *sep-1(ok1749*)/hT2g hermaphrodites with indicated transgene at 20°C.

We next attempted to examine genetic interactions with separase mutants using males in a cross to introduce the transgene. Our first attempts involved crossing transgenic males to *sep-1(e2406)*/hT2g or *sep-1(ok1749)*/hT2g hermaphrodites that also had the *unc-119* mutation to easily identify transgenic progeny. This crossing scheme should allow us to isolate F1 worms that are heterozygous mutant and heterozygous for the transgene (trans-heterozygotes, see Supplemental Figure 2A). The resulting F2 from parental crosses with SEP-1^WT^::GFP males had germline GFP expression and no lethality, but F2 animals from original parental crosses with SEP-1^PD^::GFP males lacked GFP expression and progeny did not show lethality (Supplemental Figure 2B and 2C). We had the same result when males carrying both N-terminal and C-terminal GFP fusions to SEP-1^PD^ were used (not shown). We confirmed that the hT2g balancer behaved as expected because crossing N2 males to *sep-1(e2406 or ok1749*)/hT2g hermaphrodites produced expected levels of embryonic lethality in the F1 brood (not shown). The reason for the lack of GFP expression in the F2 is not clear. One explanation is that the singled F1 worms may be non-GFP pharynx animals with WT separase from hT2g balancer breakdown, which has been previously reported (Mckim and Rose 1990). Another possibility is that the SEP-1^PD^::GFP transgene is silenced in the presence of *sep-1* mutant alleles, however we did not have this problem when crossing to homozygous mutants without hT2g (see below). In summary, we were unable to analyze genetic interactions using a scheme that started with crossing transgenic males to *sep-1(mutant)/hT2g* hermaphrodites. However, this strategy may work for other mutant and balancer chromosome combinations.

To test genetic interactions using our transgenic male propagation method, and to avoid the complications we observed with hT2 g balanced animals, we examined the phenotype of *sep-1(e2406*)/+ embryos expressing SEP-1^WT^::GFP or SEP-1^PD^::GFP. Transgenic SEP-1^WT^::GFP or SEP-1^PD^::GFP males were crossed with *sep-1(e2406*) homozygous hermaphrodites at 16°C, and F1 progeny were grown to L4 at 16°C (Figure 5A). The F1 *sep-1(e2406*)/+; SEP-1^WT^::GFP/− and *sep-1(e2406*)/+; SEP-1^PD^::GFP/− worms were shifted to 20°C at the L4 stage and embryonic lethality was determined in the F2 brood. GFP expression was confirmed in the F1 hermaphrodites used in this analysis. F2 embryos from *sep-1(e2406*)/+; SEP-1^WT^::GFP/− animals were fully viable as expected, but *sep-1(e2406*)/+; SEP-1^PD^::GFP/− F2 progeny showed 100% embryonic lethality (Figure 5B). This further demonstrates the dominant negative activity of SEP-1^PD^::GFP, because SEP-1^PD^::GFP causes lethality in separase wild-type background (Figures 2 and 3), and is further exacerbated in the presence of a mutant separase allele (Figure 4 and 5). Unfortunately all other mutant alleles of separase, including *sep-1(ok1749*), are not viable as homozygotes and could not be tested this way. These studies outline methods that can be used to examine interactions of a variety of mutant alleles and transgenes of interest by other investigators.

### Chromosome dynamics in embryos expressing SEP-1^PD^::GFP

In order to more carefully examine the dominant-negative effects of SEP-1^PD^ on chromosome segregation, we looked at phenotypes in embryos that also expressed H2B::Cherry. SEP-1^PD^::GFP; H2B::Cherry expressing worms removed from *gfp* RNAi for five generations were shifted to 25°C for 24 hours and F6 embryos (from dissected F5 adults) were imaged by confocal microscopy. We found that SEP-1^PD^::GFP followed localization patterns similar to SEP-1^WT^::GFP during the first mitosis (Figure 6A, Supplemental Movies 1 and 2). However, chromosome separation during anaphase was delayed in the presence of SEP-1^PD^ compared to SEP-1^WT^ (Figure 6B). Mitotic exit did not appear to be altered, as timing for re-condensation of chromosomes in preparation for the second division was not affected (Figure 6C). Additionally, SEP-1^PD^ expressing embryos often displayed lagging chromosomes and/or chromosome bridges. Bridges and lagging chromosomes were observed in embryos expressing SEP-1^PD^::GFP alone when separase is localized to chromosomes in anaphase, when “cross-eyed” phenotype is observed in dividing cells, and when separase is recruited to the chromosomes for the next division (n = 16/27 or 59% of embryos examined in the first and second division, not shown). A similar frequency of chromosome bridges and lagging chromosomes is observed in SEP-1^PD^::GFP embryos also co-expressing H2B::mCherry (n = 4/7 or 57% in the first and second division, Figure 6D). Chromosome bridges and lagging chromosomes were not observed in SEP-1^WT^ embryos (n= 0/19, embryos examined expressing SEP-1^WT^::GFP with or without H2B::mCherry in the first and second division, Figure 6D). These chromosome bridges and lagging chromosomes are likely due to incomplete cohesin cleavage caused by SEP-1^PD^ interfering with cohesin cleavage by endogenous wild-type separase.

**Figure 5.**
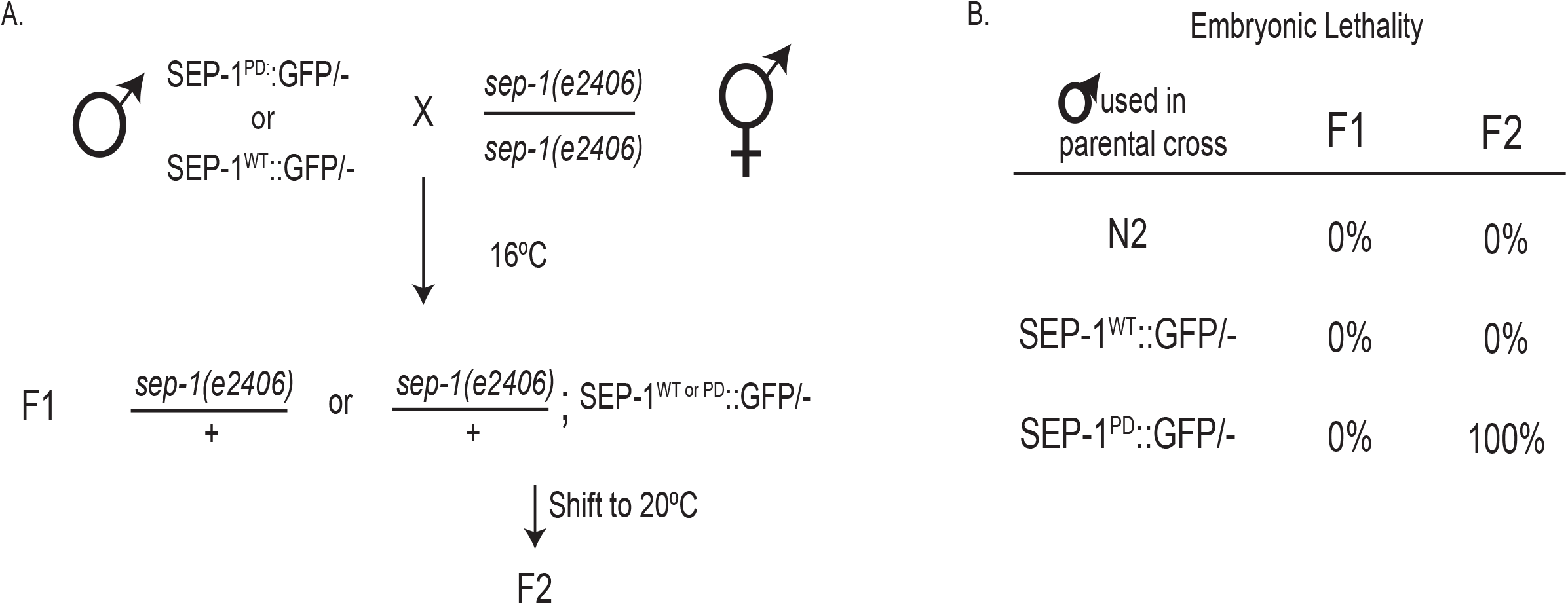
Genetic interactions of wild-type or protease-dead SEP-1 with *sep-1*(*e2406*). A. Crossing scheme of heterozygous SEP-1^WT^::GFP or SEP-1^PD^::GFP transgenic males to *sep-1(e2406)* homozygous hermaphrodites. GFP transgene expression in the F1 was determined by microscopy after the shift to 20°C. Only progeny from animals expressing SEP-1^WT^::GFP or SEP-1^PD^::GFP were analyzed. B. The table shows embryonic lethality in the F1 and F2 progeny when males carrying the indicated.

**Figure 6.**
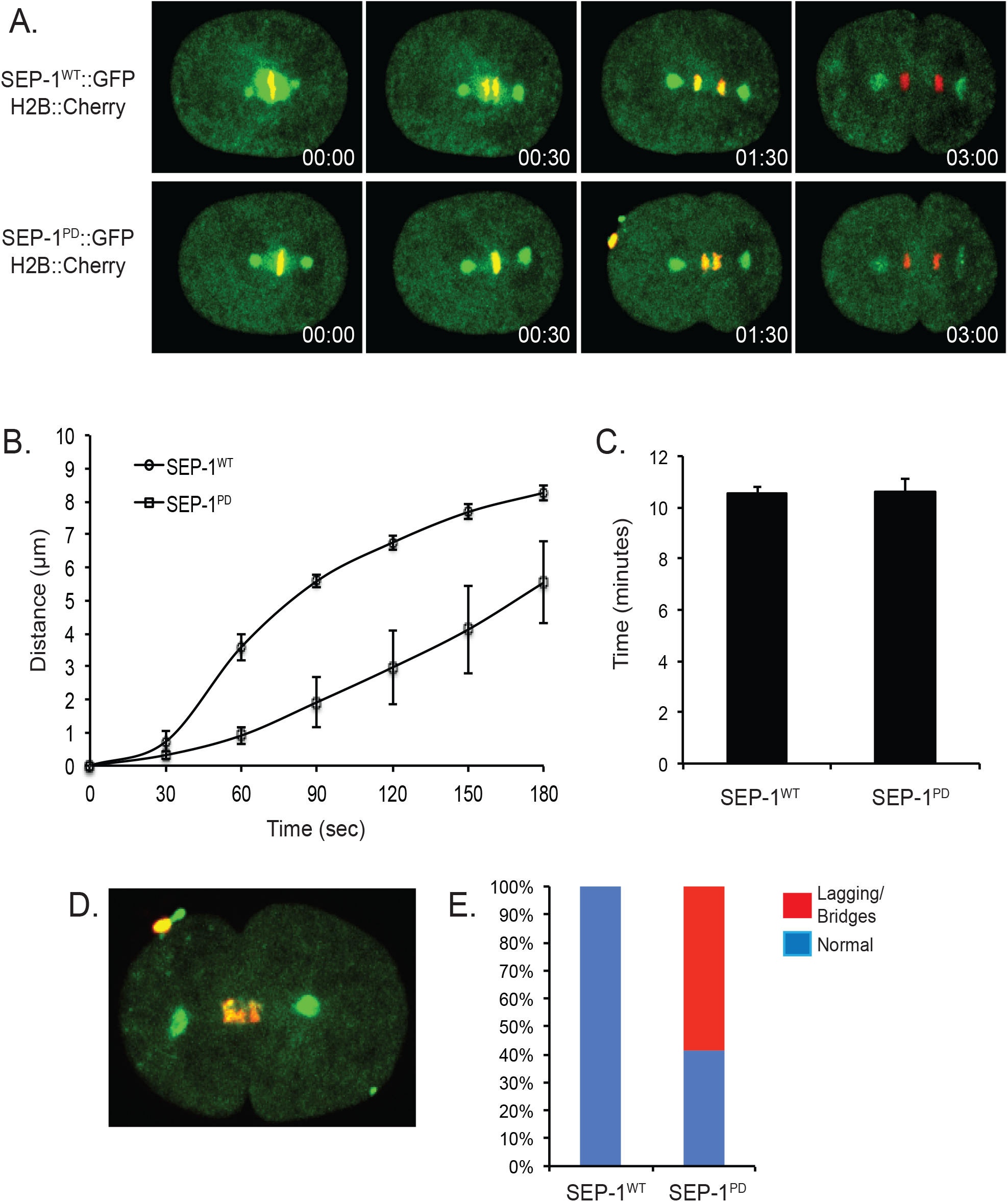
Chromosome segregation during mitosis in SEP-1^WT^::GFP or SEP-1^PD^::GFP embryos. Prior to imaging, L4 worms were shifted to 25°C for 24-48 hours. A. Live cell imaging of the first mitotic division in embryos expressing H2B::mCherry (red) and SEP-1^WT^::GFP (green, top row) or SEP-1^PD^::GFP (green, bottom row). Overlap of mCherry and GFP signal is yellow. Time stamps (minute:second) indicate the time after chromosomes move poleward in anaphase. B. Distance between chromosomes as they move poleward during anaphase of the first mitosis. Each data point represents the average distance +/− SEM, n = 10 SEP-1^PD^::GFP embryos, n = 8 SEP-1^WT^::GFP embryos. C. The average time (+/−SEM) until mitotic exit for SEP-1^WT^::GFP; H2B::mCherry (n = 8) or SEP-1^PD^::GFP; H2B::mCherry (n = 10) expressing embryos from metaphase to chromosome condensation of the following cell cycle. D. Representative image of a SEP-1^PD^::GFP; H2B::Cherry expressing embryo with lagging chromosomes during the first mitotic anaphase. E. The graph shows the percentage of embryos displaying normal chromosome separation (blue) or lagging chromosomes/chromosome bridges (red) during the first mitosis in embryos expressing either SEP-1^WT^::GFP (n = 19) or SEP-1^PD^::GFP (n = 27).

### Cytokinesis phenotype of SEP-1^PD^::GFP expressing embryos

Separase plays a crucial role in membrane trafficking during cytokinesis, and is required for cytokinesis completion (Bembenek *et al.* 2010). In order to examine whether the protease activity of separase is required for cytokinesis, we examined embryos expressing SEP-1^PD^::GFP by live imaging. Interestingly, we observed a fraction of SEP-1^PD^::GFP embryos with multipolar spindles in the one cell through four cell stage (n = 4/34 embryos examined, Figure 7A), which could indicate failed cytokinesis in the previous cell division. Since we did not observe the prior cell division in all cases where multipolar divisions were observed, we cannot rule out that these multipolar spindles might also arise from abnormal centrosome duplication, which is another known function of separase (Schvarzstein *et al.* 2013). We also observed that SEP-1^PD^::GFP expressing embryos often displayed abnormal nuclear shape in at least one daughter cell following the first or second division (n = 27/34, Figure 7C, notice in Supplemental Movie 2), that was not seen in embryos expressing SEP-1^WT^::GFP (n = 0/19 embryos). It is possible that the observed abnormal nuclear shape following division arises due to chromosome bridges and/or lagging chromosomes, as we observed that all embryos with visible lagging chromosomes and/or chromosome bridges also displayed this abnormal nuclear shape (data not shown). However, this phenotype is also observed in *syx-4* depleted embryos (Jantsch-Plunger and Glotzer 1999), and *syx-4* depletion also has a similar affect on Rab-11 trafficking during cytokinesis as separase depletion (Bembenek *et al.* 2010). These findings suggest that substrate cleavage by separase may be an essential function of separase in its various cellular roles.

**Figure 7.**
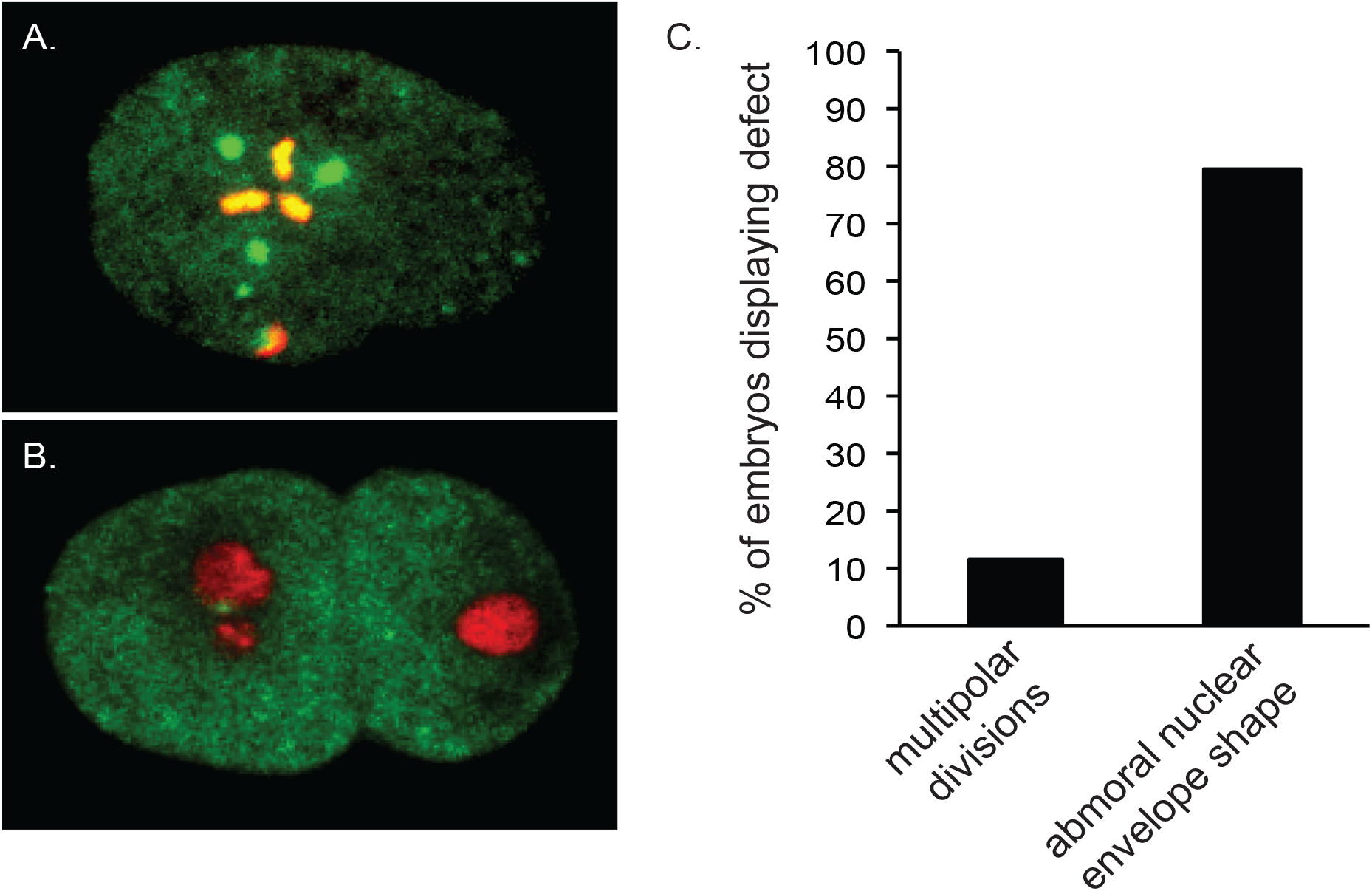
Cytokinesis phenotype of SEP-1^PD^::GFP expressing embryos. Representative phenotypes in embryos expressing SEP-1^PD^::GFP alone or with H2B::mCherry.. Phenotypes were seen in embryos with SEP-1^PD^::GFP regardless of the presence of H2B:Cherry. A. Multipolar spindle in the first division of an SEP-1^PD^::GFP; H2B::Cherry embryo. B. Abnormal nuclear envelope shape following the first mitotic division in a SEP-1^PD^::GFP; H2B::Cherry embryo. C. Percentage of SEP-1^PD^::GFP embryos that display the indicated defect. A total of n = 34 SEP-1^PD^::GFP expressing embryos were examined that express SEP-1^PD^::GFP alone or in combination with H2B::Cherry. None of these defects were observed in embryos expressing SEP-1^WT^::GFP (n = 19), not shown.

## DISCUSSION

The lack of an inducible gene expression system in *C. elegans* makes it difficult to study dominant-negative or otherwise toxic mutant proteins. Here we discuss methods that facilitate studies involving worm lines with toxic *pie-1* driven transgenes. These methods use standard laboratory techniques and are therefore feasibly implemented. While we were able to create SEP-1^PD^ transgenic lines by normal methods, recovery of bombarded animals directly on *gfp* RNAi could allow isolation of worm lines with transgenes that are more toxic than SEP-1^PD^ or allow for the isolation of overexpressing lines. These methods could also be combined with temperature sensitive mutations in the RNAi machinery to more rapidly shut off the multigenerational RNAi silencing mechanism and more quickly induce transgene expression (Calixto *et al.* 2010). In addition, bombardment of *him;unc* lines could allow for immediate isolation of transgenic males, which can be maintained by backcrossing. The male propagation method bypasses the multigenerational propagation of *gfp* RNAi, but only introduces a single copy of the transgene, which may not lead to highest expression levels. In addition, backcrossing to *unc-119* each generation can help reduce selective pressure that might silence transgene expression or select for suppressor mutations.

The *gfp* RNAi feeding and male propagation methods allow for crossing schemes to study genetic interactions of mutant proteins. Mutant separase alleles are lethal and must be maintained as heterozygotes with chromosomal balancers, which gave unexpected results with SEP-1^PD^::GFP transgenic animals. However, genetic analysis with other mutants and/or balancer chromosomes may not result in such issues and the strategies discussed above should provide guidelines to examine genetic interactions for the gene of interest. Further, male propagation and *gfp* RNAi feeding allow for the creation of double transgenic lines to study localization patterns and phenotypes by live imaging.

We employed the methods described in this manuscript to examine the consequence of SEP-1^PD^ expression in the *C. elegans* embryo. We find that protease-dead separase is dominant negative and interferes with normal separase function in our system. A dominant negative activity of protease-dead separase has not been reported in other systems. This dominant negative activity could arise due to substrate trapping by the mutant separase enzyme, preventing substrate cleavage by endogenous separase and therefore interfering with essential separase function and leading to death of the embryo. Consistent with this, we found that chromosome separation during anaphase is delayed and there is a high frequency of lagging chromosomes and chromosome bridges in embryos expressing SEP-1^PD^ compared to SEP-1^WT^, suggesting that cohesin cleavage is incomplete in the presence of SEP-1^PD^.

Whether substrate cleavage by separase underlies its role in vesicle trafficking is not known. Although we did not see high levels of cytokinesis failure or indications of eggshell permeability in transgenic embryos expressing SEP-1PD in the *unc-119* background, endogenous separase was present and could have enough activity for sufficient exocytosis to support these events. Additionally, exocytosis could be slowed or delayed, yet occur at sufficient levels to avoid more severe phenotypes. Indeed, RAB-11 trafficking is affected in embryos depleted of separase by RNAi, even if they do not fail cytokinesis (Bembenek *et al.* 2010). Unfortunately we were unable to make a line expressing both Cherry::RAB-11 and SEP-1^PD^::GFP despite several attempts. Further analysis is required since previous work indicated that polar body extrusion is independent of separase’s proteolytic activity in mouse oocytes (Kudo *et al.* 2006). However, this analysis was performed in separase-null mouse embryos, which may have a different phenotype than seen in wild-type separase background. For example, separase could require autocleavage to efficiently bind substrates, which could be mediated in our SEP-1^PD^ transgenic lines by endogenous separase. Further analysis must be performed to more closely examine the proteolytic function of separase in exocytosis and to identify putative substrates involved in membrane trafficking.

